# 3D map-guided modeling of functional endometrial tissue using multi-compartment assembloids

**DOI:** 10.1101/2025.08.22.671545

**Authors:** Kehan A. Ren, Victoria Duarte-Alvarado, Xin Di Zhou, Bryan Ho, André Forjaz, Ashleigh J. Crawford, Gretchen M. Alicea-Rebecca, Saurabh Joshi, Praful Nair, Eban A Hanna, Denis Wirtz

**Author notes:** These authors contributed equally to this work (Kehan A. Ren, Victoria Duarte-Alvarado).

## Abstract

The human endometrium is a dynamic tissue that lines the uterus and undergoes constant remodeling, making it especially susceptible to gynecological diseases like endometriosis and endometrial cancer. The molecular mechanisms of these conditions are not well understood, partly due to the lack of in vitro models that mimic endometrial physiology, which limits options for targeted intervention and treatment of these diseases. Mouse models are also inadequate, as common laboratory strains do not naturally undergo a menstrual cycle comparable to that of humans. This study addresses this need by developing a 3D multi-compartment assembloid that mimics the architecture of endometrial tissue and recapitulates all three phases of the menstrual cycle (proliferative, secretory, and menstrual regression) within a single platform. The cellular and extracellular matrix (ECM) components in each compartment are carefully tuned based on a 3D spatial cellular map of endometrial tissue. The model contains endometrial epithelial cells enveloped in a basement membrane and endometrial stromal cells in a surrounding collagen-rich layer; this architecture allows realistic interactions between these cells and their respective ECMs. This assembloid successfully supports the controlled growth and organization of these cells, revealing reciprocal regulation of cell behavior and exhibiting compartment-specific hormonal responses, i.e., stromal decidualization. This platform enables the study of dynamic, phase-resolved, and compartment-specific paracrine signaling in human endometrial biology. By combining tissue-informed design, modular fabrication, and full-cycle hormonal responsiveness, this model sets a new benchmark for blastocyst implantation studies, organ modeling, and precision diagnostics in human reproductive health.

## Introduction

The endometrium is a critical component of the human uterus that undergoes highly dynamic, cyclical remodeling during a woman’s reproductive years in preparation for potential pregnancy (*1*). This frequent remodeling, combined with exposure to both endogenous and exogenous factors, makes endometrial tissues particularly susceptible to pathological conditions, including gynecological diseases, such as endometriosis and endometrial cancer (*2*, *3*). These diseases affect millions of women worldwide and significantly impact quality of life (*4*, *5*). Diagnosis requires surgical evaluation and histological confirmation. Current standard-of-care treatments include the use of non-steroidal pain medications to treat symptoms, hormone-based therapies, surgery, adjuvant radiotherapy, chemotherapy, and total hysterectomy with or without oophorectomy. These therapies, often met with high recurrence rates, are contraindicated in individuals seeking to preserve fertility or pursue pregnancy (*6*, *7*). A major barrier to therapeutic advancement is the incomplete understanding of the complex physiological environment of the endometrium (*8*). This limitation stems from the lack of validated models that recapitulate the complex cellular heterogeneity, architecture, mechanical functions and native cell–cell and cell–extracellular matrix (ECM) interactions of the endometrium (9–11). There is also no 3D map of the endometrium at the single cell level.

The cellular and non-cellular composition, the complex architecture, and the intercellular interactions in the endometrium are not captured in traditional 2D in vitro cultures (12). Mouse models are also inadequate, as key endometrial diseases do not naturally occur in mice (9, 10, 13). Only 3D systems composed of human cells can replicate the cellular diversity, architecture, and mechanics of the tissue (10, 11). Recent advances in 3D in vitro modeling have substantially enhanced our ability to interrogate human endometrial biology in systems that recapitulate tissue-specific structure and function (*14–37*, *37–40*). Organoid models, derived from primary endometrial epithelial fragments embedded in solubilized basement membrane (Matrigel), have been instrumental in enabling long-term culture of hormone-responsive epithelium and have yielded key insights into endometrial epithelial lineage dynamics, secretory transformation, and hormone-regulated gene expression (*41*, *42*). Nonetheless, these models only contain epithelial cells and lack stromal components (35). Their reliance on bulk Matrigel and specialized media further limits interlaboratory reproducibility and constrains the study of cell–ECM or tissue–tissue interactions.

To overcome these limitations, endometrial stromal–epithelial organoids co-culture models have emerged, where epithelial organoids are co-embedded with primary stromal cells in either collagen-based or synthetic extracellular matrices for improved anatomical accuracy (*43*) . These platforms have enabled the investigation of paracrine signaling and hormone-mediated remodeling across compartments, with recent advances employing defined, tunable ECMs to improve anatomical relevance and experimental control. However, these systems generally rely on sequential assembly within a shared ECM, where endometrial epithelial and stromal populations are matured separately before being combined, limiting opportunities for synchronized co-development and obscuring compartment-specific matrix signaling (*19*, *43*). Assembloids represent the most integrated approach to date, spatially organizing endometrial epithelium and stromal cells into physiologically relevant 3D configurations (*44*). These models have advanced functional studies of decidualization, inflammation, and embryo implantation, but often capture only discrete phases of the menstrual cycle, most commonly the proliferative or mid-secretory stages, and rarely simulate menstrual regression (*16*, *44*).

Here, we present a novel, highly tunable multi-compartment assembloid model that enables the coculture of endometrial epithelial and stromal cells within spatially distinct, tissue-specific ECM compartments (which we call a coculture assembloid), designed to replicate the native architecture and functional complexity of the human endometrium (*45*). The cell types and ECMs in each compartment are carefully selected and optimized based on histological references and informed by a new 3D map of the human endometrium generated using the AI-based spatial tissue-mapping method CODA(*45–47*). By seeding endometrial epithelial and stromal cells in anatomically relevant adjacent matrices, this model supports physiologically correct cell-cell and cell-ECM interactions that drive coordinated development.

Unlike previous co-culture or assembloid systems that rely on shared ECM environments or sequential assembly strategies, our platform enables synchronized tissue development within ECM-defined, compartment-specific niches. Structurally, molecularly, and functionally validated, the assembloids form mature epithelial networks that resemble the interlinked glandular structures observed in the CODA map of endometrial tissue, retain apicobasal polarity, and express characteristic lineage markers. The model exhibits robust responsiveness to ovarian sex hormones, including estradiol- and progesterone-driven transitions, and uniquely captures hormone withdrawal-induced menses using RU-486, allowing simulation of the entire menstrual cycle *in vitro*.

## Materials and Methods

### Experimental design

Traditional *in vitro* models of the human endometrium have been limited in their ability to recapitulate the tissue’s multicellular architecture and supportive stromal microenvironment needed for physiological function. To address these limitations, we developed a three-dimensional (3D) multi-compartment assembloid model that recapitulates human endometrial tissue structurally, molecularly, and functionally, guided by a 3D reconstructed reference map of endometrial tissue generated using AI-based 3D tissue mapping method CODA (*46–48*). We employed an oil-in-water droplet method to assemble the endometrial epithelium core and stromal corona sequentially (*45*, *49*). For validation, we assessed the assembloids for anatomical fidelity and functional competence. Immunofluorescence and Western Blot analyses were performed to verify proper glandular structure formation and the expression of characteristic epithelial and stromal markers. We further evaluated hormonal responsiveness by treating assembloids with 17β-estradiol and progesterone, confirming physiologic functional responses in both compartments.

### CODA mapping of eutopic endometrium

To generate CODA maps of eutopic endometrium, we first serially sectioned endometrial tissue samples retrieved from pathology archives at Johns Hopkins Medicine from donors with no history of disease (**Fig. 1a**). All procedures were approved by the Johns Hopkins Internal Review Board. The tissues were transported in saline on wet ice, formalin fixed (10%, VWR 16004-121), and paraffin embedded. The entire tissue was serial sectioned at 4-μm thickness at Johns Hopkins Oncology Tissue Services. Tissue sections were stained with hematoxylin and eosin (H&E). Tissue sections were scanned at 20× or 40× magnification (Hamamatsu NanoZoomer S210). One in every two tissue sections was H&E stained.

To train a deep-learning model to semantically segment these tissue components in the entire stack of tissue sections, cellular or extracellular components were manually labeled from a small subset of these H&E images. Our main tissue components of interest were identified and labelled as the glandular epithelium and endometrial stroma (**Fig. 1b**). Following the method we have devised and detailed in (*48*, *50*) to train the model and align completely labelled section, we produced 3D maps of the tissue at the single-cell level. Tissue architectural parameters were quantified with CODA maps. The number of distinct cell types per unit volume of their corresponding tissue components (e.g., the number of epithelial cells in the endometrium), the ratios of epithelial to stromal cells within a 3 mm diameter of the epithelium (distance determined by assembloid corona size), the volume of the gland and lumen, and the depth of the gland across successive sections are all examples of quantifiable tissue parameters. Additionally, the 3D reconstructions made it possible to quantify the total number of glands in a single tissue and visualize the 3D connection of glands that were visible in a 2D tissue segment.

### Cell culture

Endometrial epithelial and stromal immortalized cells (hEM3 endometrial epithelial cells and 9700N gynecological stromal cells) were obtained from Dr. Tian-Li Wang’s lab at Johns Hopkins Medicine and expanded in cell culture medium containing RPMI-1640 (Gibco 11875-093) supplemented with 10% Fetal Bovine Serum (FBS, Corning 35-010-CV) and 1% Penicillin-streptomycin (Sigma P0781) at 37°C with 5% CO_2_ in the incubator. Luciferase and mCherry were artificially expressed by hEM3 and GFP by 9700N. Endometrial Uterine primary epithelial cells (LifeLine Cell Technology FC-0078) were cultured in complete ReproLifeTM Medium (LifeLine Cell Technology LL-0068) for up to 3 passages. Normal Human Uterine Primary Fibroblasts (LifeLine Cell Technology FC-0076) were cultured in complete FibroLife Medium (LifeLine Cell Technology LL-0011) for up to 3 passages. All cells were cultured at 37°C and 5% CO_2_ in a humidified incubator and did not surpass 15 cell passages.

### Multi-compartment assembloid

#### Oil columns preparation

To prepare the oil columns, a new wafer of filterless 10 µL pipette tips was loaded into a pipette tip box (USA Scientific 1111-3700). Using a P10 multichannel pipette, 10 µL of cell culture medium was added to the bottom of each pipette tip. The tips were then carefully returned to their original positions, leaving the liquid in place. Next, a P300 multichannel pipette was used to add approximately 100 µL of mineral oil (Sigma 69794) to the upper portion of each pipette tip forming stable oil columns.

#### Fabrication of assembloid core

Endometrial epithelial cells (4 × 10^6^) were pelleted by centrifugation at 1000 RPM for 5 minutes in a 1.5 mL Eppendorf tube. The pellet was gently resuspended in 100 µL growth factor-reduced Matrigel (Corning 354230), keeping the mixture on ice to prevent mature gelation and minimizing bubble formation. For droplet loading, a 10 µL low-binding pipette tip was primed by drawing up 1 µL of mineral oil and dispensing it onto a gloved hand to coat the inner surface of the tip; any excess oil on the tip was removed. Using the primed tip, 1 µL droplets of the cell–Matrigel mixture were dispensed into individual oil columns. The tip was wiped between each drop to ensure consistency. This process was repeated, changing tips every 1-2 columns of the tip box, with each tip primed. The embedded cores were incubated at 37 °C and 5% CO₂ for 1 h to allow for gelation. Meanwhile, 10 mL of culture medium was pre-warmed in a 37 °C water bath for downstream use.

#### Core harvest and wash

After gelation, the cores were collected from the oil columns using a cut 300 µL pipette tip and transferred into a 15 mL conical tube containing 4–7 mL of pre-warmed culture medium. Any mineral oil that came in the transfer was carefully vacuumed from the top. Some medium was also aspirated, being careful not to disturb the cores, until only 2-3 ml of core-containing medium was left in the conical tube so the cores could be transferred to a new 15 ml conical tube using a cut 1000 µL pipette tip without getting the pipette inside the liquid. This step was repeated several times to ensure no oil stayed with the cores, as this would make the core separate from the corona in the future.

#### Encapsulation of assembloid corona

To prepare our collagen layer, 500 µL of collagen I solution was prepared at the desired concentration. If stromal cells were to be included, they were suspended thoroughly into the collagen solution to ensure homogenous cell density. Approximately 20 cores were transferred into 50 µL of the collagen mixture using a cut 300 µL pipette tip, then gently mixed. Using a cut P20 pipette tip single cores were individually aspirated in 10 µL of collagen, ensuring the core was centered in the collagen droplet. Each droplet was then dispensed into fresh oil columns. Each pipette tip was cleaned between droplets to avoid oil disturbing the adequate formation of the assembloids. All assembloids were incubated at 37 °C and 5% CO₂ for 1.5 h to complete gelation of collagen. In parallel, 15 mL of cell culture medium was pre-warmed.

#### Assembloid harvest and wash

Fully gelled assembloids were harvested from the oil columns using a cut 300 µL pipette tip and placed into a 15 mL conical tube containing 4-7 mL of warm medium. Any remaining oil and excess medium were aspirated as in the previous wash. To ensure complete cleaning, assembloids were transferred into a new 15 mL conical tube using a cut 1000 µL pipette tip and complete cell culture medium was added for plating.

### Plating and culture

Individual assembloids were plated into 96-well round-bottom plates (Genesee Scientific 25-224), using a cut 200 µL pipette tip in 200 µL of medium. one assembloid was placed per well. The assembloids were cultured at37 °C in a humidified incubator with 5% CO₂ and was medium changed every 2-3 days until samples were collected for downstream applications.

### Standard organoid

The protocol for the standard organoid generation was previously described (*41*). Briefly, Endometrial epithelial cell pellets were resuspended in growth factor reduced Matrigel with pellet volume: Matrigel volume at 1: 5. A 20 µL drop of Matrigel-cell mixture was then plated on a glass bottom plate and gelled for 1 h. The organoid was maintained in the organoid expansion medium at 37°C with 5% CO_2_ in the incubator, as described (*41*). Media changes were every 3 days.

### PrestoBlue proliferation assay

PrestoBlue cell viability assay was performed on the assembloid to track epithelial proliferation. At first, 100 µL of medium was removed from every well in the assembloid culture plate, and 100 µL of medium was added to 3 -5 blank wells as a background reference. 10X PrestoBlue reagents (ThermoFisher A13262) were diluted to 2X in cell culture medium. 100 µL of 2X PrestoBlue solution was then added into every well to dilute PrestoBlue to 1X final concentration. Then the assembloids were incubated avoiding light at 37 °C with 5% CO_2_ for 3 h. The red fluorescence (RFU) was read on a SpectraMax plate reader. After reading, assembloids were washed with DPBS (Corning 21-031-CV) thrice and washed with cell culture medium thrice, then replenished with 100 µL of medium. The plate was put back into the incubator and cultured until the next timepoint.

### Luciferase assay

Luciferase assay was used as an endpoint assay to measure Luciferase-tagged epithelial cell proliferation in the coculture with stromal cells. Dead control wells were prepared as the reference of background signals one day ahead of the measuring day. To prepare dead control wells, 180 µL of an assembloid with medium was transferred to the white microplate, and 20 µL of 10% Triton-X solution (Sigma T9284) was added to each well to kill the cells. On the measuring day, 4 - 5 assembloids were washed with DPBS thrice, and 50 µL of the assembloid with medium was transferred to the same white microplate. 50 µL of 4 mg/mL collagenase (Gibco 17018029) solution was added to each well. Then the plate was put into the incubator. After 1 h, 100 µL of Luciferase reagent was added to each well. The plate was then shaken at 200 rpm for 5 minutes and had the luminescence reading on a SpectraMax plate reader.

### Hormone treatment

The hormone treatment timeline is described in (**Fig. 5a**). The generated assembloids were maintained in Phenol red-free RPMI-1640 (Gibco 11835030) supplemented with 10% Fetal Bovine Serum and 1% Penicillin-streptomycin at 37°C with 5% CO_2_ in the incubator until day 3. Hormones were pre-diluted in ethanol prior to dilution to the phenol red-free medium. At day 3, 100 µL of medium was substituted with fresh medium containing 40 nM of E2 for the proliferative phase induction, and the final E2 concentration for cell culture was 20 nM. Secretory phase induction started at day 7, 100 µL of medium was substituted with fresh medium containing 2 µM of P4 and 40 nM of E2. At day 12, E2 and P4 were washed out from the culture medim, and 1 µM of RU-486 was added to the fresh medium, aiming to induce the menstrual phase.

### Assembloid imaging

Assembloids were imaged on a Nikon Ti2 microscope (phase-contrast) and a Nikon A1R confocal/Nikon Eclipse Ti inverted microscope (DIC, fluorescence and immunofluorescence confocal). Assembloids were collected in the 1.5 mL Eppendorf tube and washed once with DPBS. The assembloids were fixed in 4% paraformaldehyde (PFA) for at least 1 h at 4 °C. After that, cells fixed in the ECM were permeabilized with 0.5% Triton-X-100 (Sigma T9284) diluted in DPBS for 1 h at room temperature and blocked with 5% Normal Goat serum (NGS) for 3 h. Primary antibodies used for IF are listed in **Table 1**. Primary antibodies were diluted in 1% NGS using the pre-decided dilution rate and added to the assembloid for overnight incubation at 4 °C. Antibody diluent was removed and washed with DPBS three times. Secondary antibodies were diluted in 1% NGS at 1:100 along with 1:200 Hoechst 3342 (ThermoFisher H3570) and 1:400 phalloidin conjugates (Invitrogen A12379) and incubated with assembloid for 3 h at room temperature. After 3 washes with DPBS, fructose-glycerol clearing solution (60% glycerol, 2.5 M fructose in DI water) was added to cover the assembloids and incubate for 20 minutes at room temperature to enhance signal quality (*51*). To image the stained assembloids with a confocal microscope, an assembloid with 50 µL DPBS was placed on a petri dish with a glass bottom. For the immunofluorescent images, maximum intensity projection of z-stacking images was processed by ImageJ (FIJI).

### Western blot assay

Protein samples were prepared by lysing the assembloids in 2x clear sample buffer (120 mM Tris pH 6.8, 4% SDS). Lysates were sonicated with a needle probe sonicator (15 pulses, VWR Scientific), denatured at 100°C for 5 minutes and centrifuged at 16,000 rpm for 15 min at 4 °C. Supernatants were collected and stored at -80°C, and protein concentration was determined using a microBCA protein assay (ThermoFisher, 23235). Equal amounts of protein (45 µg per lane) were mixed with 2× Laemmli sample buffer (BioRad 1610737) containing 10% β-mercaptoethanol and heated at 100 °C for 5 min, then microcentrifuged for 30 seconds. Proteins were separated by 4-15% precast polyacrylamide gels (BioRad PowerPac) at 165 V for 45 min and transferred onto 0.2 µm PVDF precut blotting membranes (Bio-Rad) using a Trans-Blot Turbo transfer system (BioRad 1704156) in 7 minutes. Membranes were blocked in 5% non-fat milk in Tris-buffered saline with 0.1% Tween-20 (TBST) for 1 h at room temperature. Blocking was followed by a 5-minute wash in TBST and membranes were incubated overnight at 4 °C with primary antibodies diluted in blocking buffer. After three washes with TBST (5 min each), membranes were incubated with HRP-conjugated secondary antibodies anti-rabbit (7074) and anti-mouse (7076) also diluted in 5% nonfat milk in TBST at 1:2000 ratio for 1 h at room temperature. Signal detection was performed using Prometheus Femto ECL Reagent (Genesee Scientific 20-302) and visualized on a ChemiDoc XRS+ Imaging system (Bio-Rad). Band intensities were quantified using Image Lab (Bio-Rad) software and normalized to housekeeping controls (e.g., HSP90 or GAPDH) as indicated.

### RTqPCR assay

Primers were ordered from IDT and reconstituted at 100 µM. Total RNA from the samples was first extracted and quantified using a nanodrop. RNA was then converted to cDNA to amplify the level of gene expression. RT-qPCR reactions were prepared using a SYBR Green-based detection system. Each 10 μL of RT-qPCR reaction contained 5 μL of SYBR Green Master Mix, 1.5 μL of cDNA, 3 μL of qPCR water and 0.5 μL of a master mix containg qPCR water, forward and reverse primers (All primer sequences are listed in **Table 2**). Reactions were set up in triplicates in a 384-well plate, with a no-template control included to check for contamination. RT-qPCR was performed on a CFX384 Real-time system (Bio-Rad) using the following thermal cycling conditions: Initial denaturation at 95 °C for 30 s, followed by 40 cycles of 95 °C for 10 s and 60 °C for 30 s. Real time data was imported to the CFX Manager software (Bio-Rad) and the quantification cycles (Cq) normalized to a panel of housekeeping genes that consists of B-Actin, GAPDH, aTub, 18s, RPL13A. All analyses were performed using data from six biological replicates, each with 3 technical triplicates.

**Table 1.**
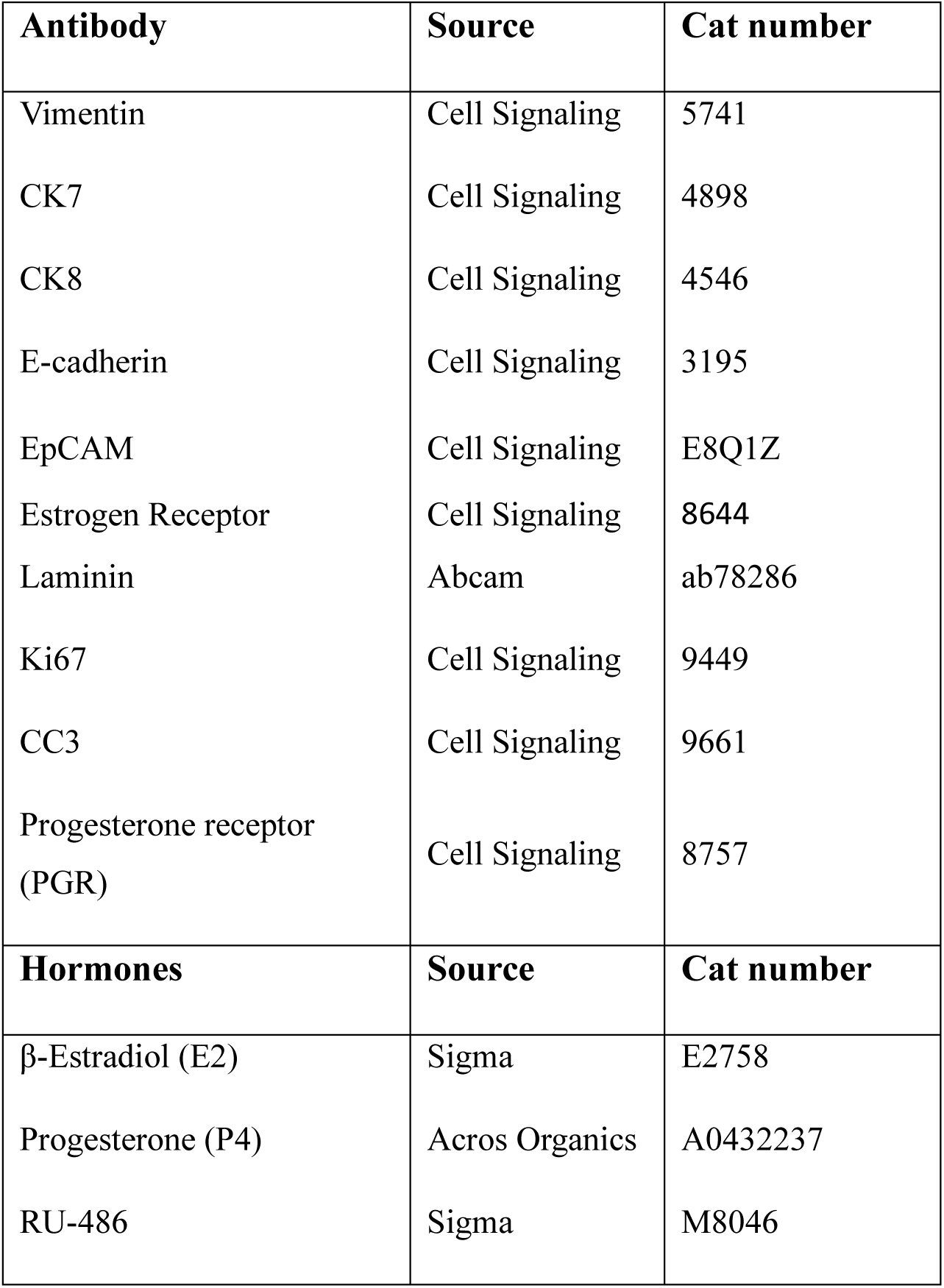
Antibodies and hormones used.

| Antibody | Source | Cat number |
| --- | --- | --- |
| Vimentin | Cell Signaling | 5741 |
| CK7 | Cell Signaling | 4898 |
| CK8 | Cell Signaling | 4546 |
| E-cadherin | Cell Signaling | 3195 |
| EpCAM | Cell Signaling | E8Q1Z |
| Estrogen Receptor | Cell Signaling | 8644 |
| Laminin | Abcam | ab78286 |
| Ki67 | Cell Signaling | 9449 |
| CC3 | Cell Signaling | 9661 |
| Progesterone receptor (PGR) | Cell Signaling | 8757 |
| Hormones | Source | Cat number |
| β-Estradiol (E2) | Sigma | E2758 |
| Progesterone (P4) | Acros Organics | A0432237 |
| RU-486 | Sigma | M8046 |

**Table 2.**
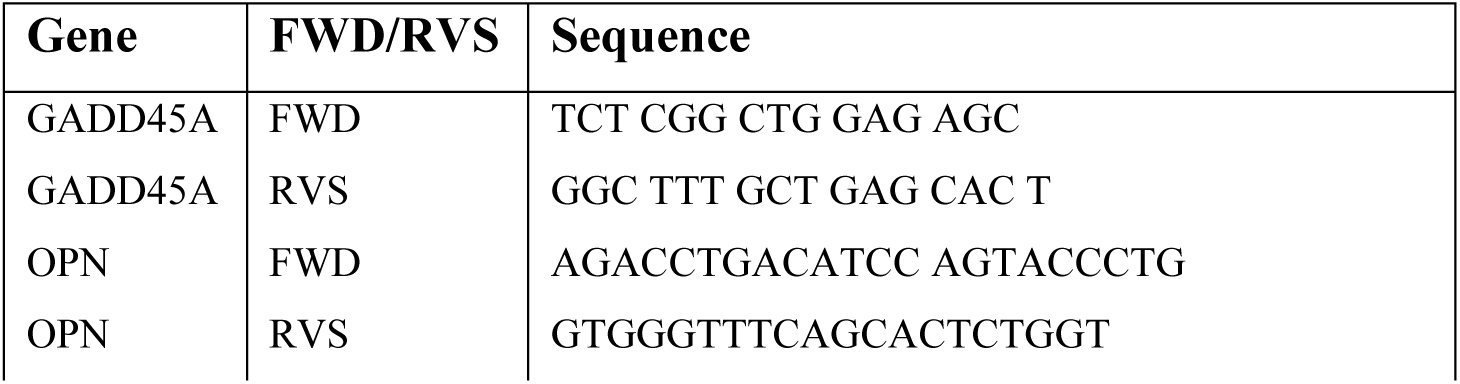
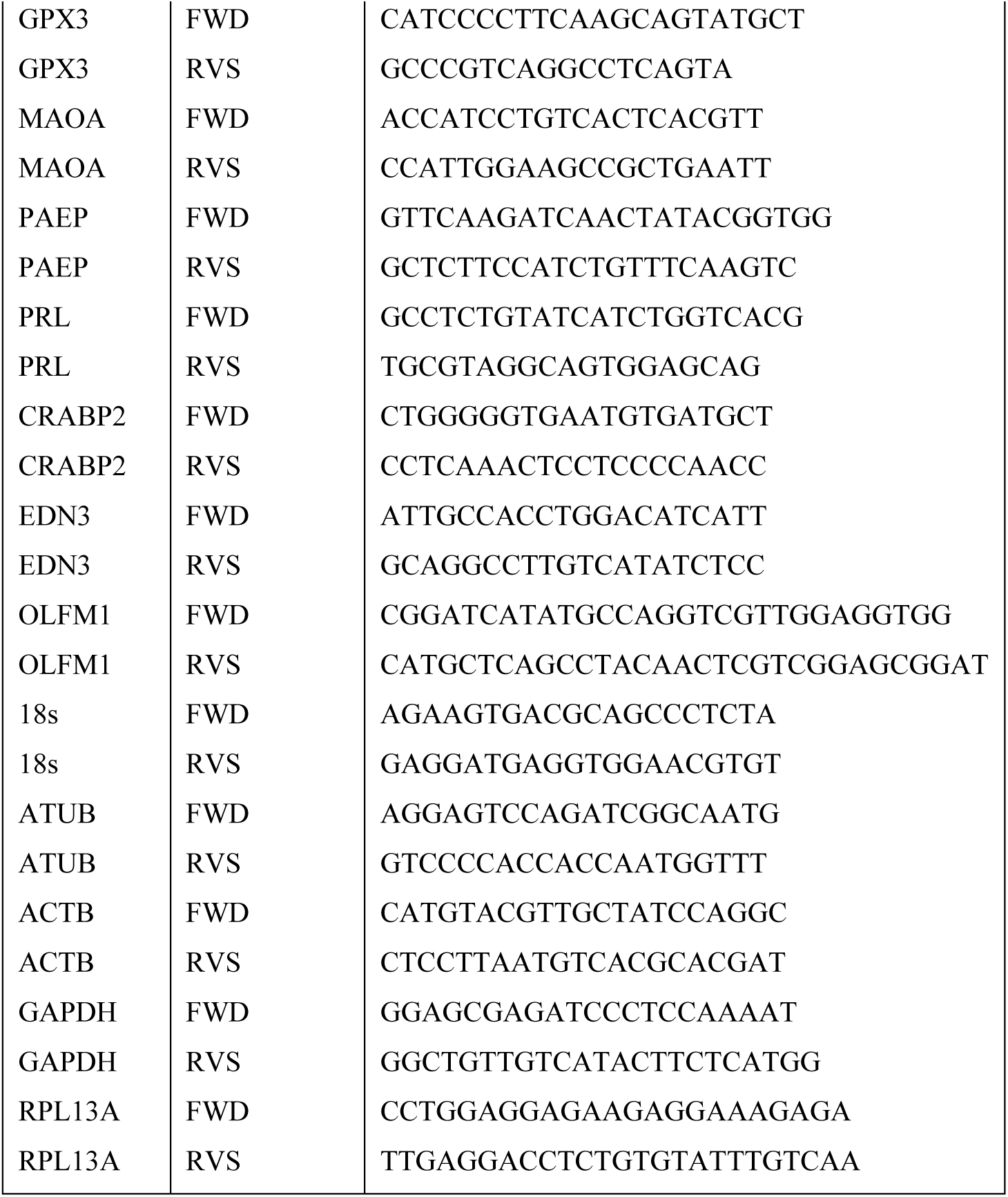
Primers used in RT-qPCR.

## Results

### The multi-compartment assembloid is designed to mimic the microenvironment of the human endometrium

We developed a multi-compartment assembloid model to produce a structurally, molecularly, and functionally accurate model of the human endometrium. In particular, it is designed to mimic the microenvironment where epithelial glands are supported by a basement membrane and surrounded by densely packed, collagen-rich endometrial stroma (**Fig. 1a-b**). Human endometrial epithelial cells (either cultured or primary) were resuspended in a droplet of Matrigel (i.e., the assembloid core), and the endometrial stromal cells (either cultured or primary) were resuspended in a surrounding collagen I corona (**Fig. 2a**). This biphasic configuration contrasts with the conventional organoid model as described above (*41*), where endometrial epithelial cells are cultured in a solubilized basement membrane, such that a single cell type is cultured in a single ECM compartment (**Fig. 2b**). The core and corona were generated on the same day to allow stromal regulation of epithelium formation starting from the beginning of endometrium regeneration (*52*).

**Figure 1.**
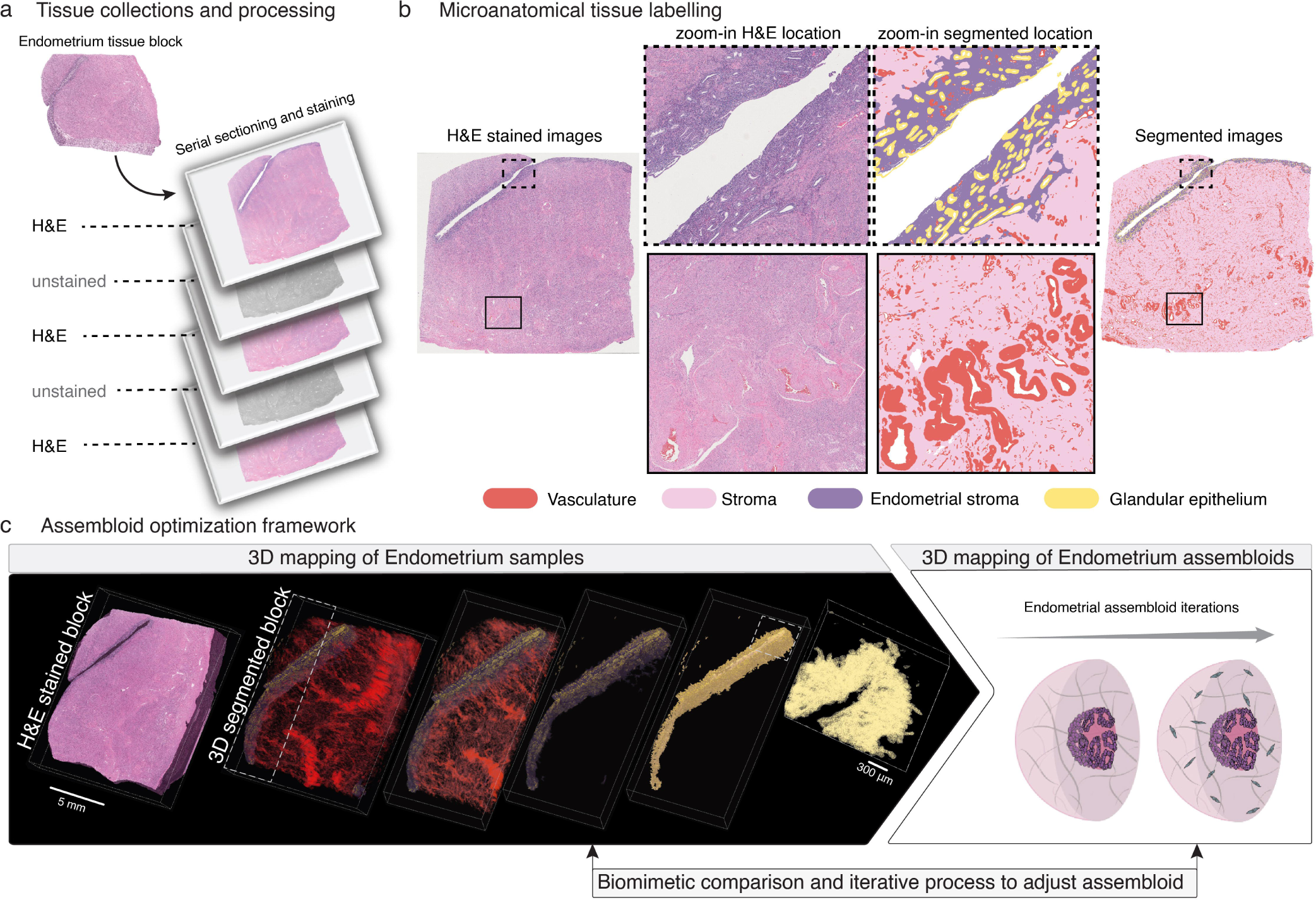
Spatial architecture of the human endometrium revealed by CODA-based 3D tissue mapping. (**A**) A human eutopic endometrium obtained from donors with no history of disease was formalin fixed, paraffin embedded and serially sectioned at 4-μm thickness. Every other section was stained with hematoxylin and eosin (H&E) and scanned at 20× or 40× magnification. (**B**) A deep-learning model was trained to semantically segment tissue components using manually labeled H&E images from a subset of sections. The primary components of interest were the glandular epithelium and endometrial stroma. (**C**) 3D reference map of human eutopic endometrium generated using the computational platform CODA. The model was trained and applied to the aligned sections to generate single-cell resolution 3D reconstructions. Endometrial stroma (purple) and glandular epithelium (yellow) are isolated for visualization. A single-cell resolution zoom in of glandular epithelium shows that epithelial ducts are interconnected in 3D.

To guide our initial cell seeding density and ECM parameters, we used the 3D single-cell resolution reference map of a non-diseased human endometrial tissue generated using CODA (**Fig. 1**). To inform our custom endometrial assembloid design and verify the resulting assembloid architecture, an archived endometrial tissue pathology sample was serially sectioned, stained, and scanned at sub-micron resolution (**Fig. 1a**). The images were registered with sub-micron resolution and a convolutional neural networks (CNN) algorithm was trained and validated to detect and segment the glandular epithelium, endometrial stroma, myometrium, and vasculature. Hematoxylin and eosin channels were deconvolved to identify individual nuclei in these tissue components. This map revealed a dense packing of stromal cells surrounding a complex network of interlinked glandular epithelium, which we aimed to capture in our assembloid model.

The first parameter that we adjusted for the design of the assembloid was the concentration of collagen I in the stromal compartment, which was increased to improve the structural stability of the outer collagen-rich corona, preventing collapse during long-term culture. This modification also enhanced anatomical fidelity by better mimicking the compact, ECM-dense stroma seen in H&E-stained sections of native endometrial tissue, particularly in terms of collagen fiber density and spatial organization. To determine the right concentration of collagen I, we tracked the assembloid development via phase-contrast microscopy. The final concentration of collagen I chosen for the assembloid design was 4 mg/mL; lower concentrations did not support long-term culture (**Fig. 2c**). In the endometrial tissue, we observed dense stromal cell packing (**Fig. 1a**), which is another feature that could be mimicked in the assembloid design (**Fig. 2d**). Stromal cells seeded in the collagen I were previously shown to remodel the ECM (*53*, *54*), and we observed that stromal cells induced changes in the corona thickness and the overall morphology of our assembloids (**Fig. 2e**). Furthermore, by tagging our epithelial cells with luciferase, we could measure their relative cell count without interference from the stromal cells. As expected, the epithelial proliferation was significantly suppressed when the stromal cells were added to the conditioned medium and to the collagen corona (**Fig. 2f**). For the initial stromal cell seeding density, we tested 200, 500, and 1 × 10^3^ stromal cells/μL collagen. By day 6, only the 1 × 10^3^ cells/μL condition consistently resulted in dense stromal cell networking, suggesting that this concentration provides the most effective parameter for establishing a physiologically relevant stromal compartment (**Fig. 3a**).

Epithelial morphology is another important factor that was considered carefully. The CODA map showed that the glands were more intricately linked and complex than observed in standard 2D histological tissue sections (**Fig. 1b**). In the first generation of assembloids, we seeded 1 × 10^4^ cells/µL epithelial cells in the Matrigel core (**Fig. 2e**). When the epithelial cells were cultured in the Matrigel core with an empty collagen corona (which we call monoculture assembloid), we were able to identify a portion of the interconnected epithelium by monitoring the growth of the endometrial structure. As seen in **Fig. 2f**, and as previously observed by (*55*), stromal cells regulate epithelial growth, which made epithelium rapidly lose its networking structure once they were introduced to the collagen layer. As we increased the initial epithelial cells to 4 × 10^4^ cells/uL in each Matrigel core, the epithelial network started to form in as little as 3 days and matured by day 9, with more connections between the epithelial branches (**Fig. 3b**). When the epithelial cells were cocultured with stromal cells, the interconnection of the epithelium persisted and was more complex than the monoculture system.

These optimizations established a structurally stable and physiologically relevant multicompartment assembloid that recapitulates key architectural and functional features of the human endometrium, providing a robust platform for modeling dynamic stromal-epithelial interactions during the menstrual cycle.

**Figure 2.**
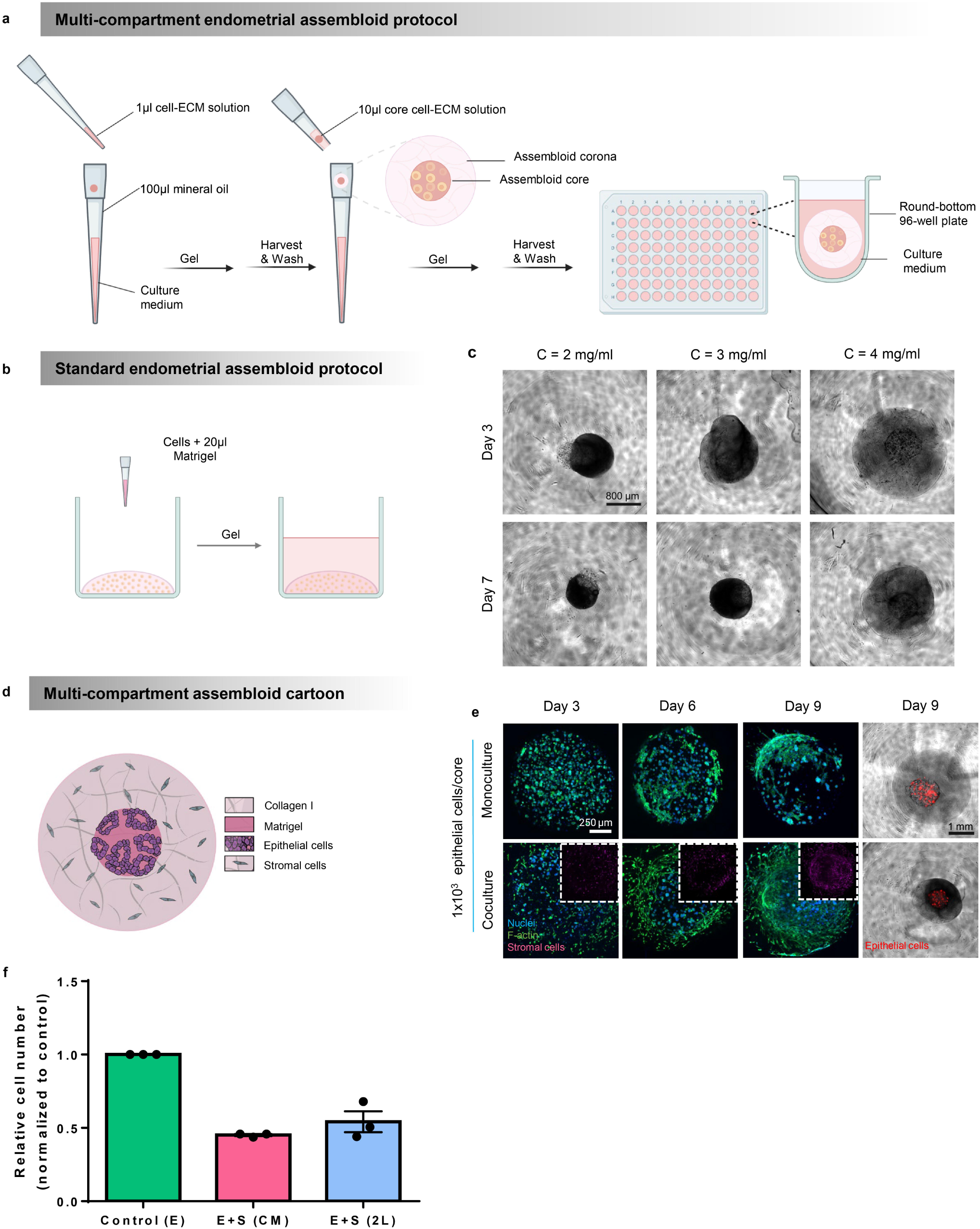
Design of endometrial multi-compartment assembloids. (**A and B**) Schematics to generate multi-compartment assembloids (**A**) and standard endometrial organoids (**B**). (**C**) Phase-contrast images of endometrial epithelial cells cocultured with gynecologic stromal cells in the corona with increasing collagen concentration. 4X magnification. Scale bar, 800 µm. (**D**) Cartoon depicting the assembloid design with coculture of epithelial and stromal cells in distinct ECM compartments. (**E**) Confocal microscopy images of endometrial epithelial cells cocultured with gynecologic stromal cells in the corona with increasing stromal cell concentration. Magenta, stromal cells. Blue, nuclei. Green, F-actin. 10X magnification. Scale bar, 250 µm. (**F**) Relative proliferation of epithelial cells in multi-compartment assembloids measured by a luciferase assay. Assembloid conditions include Control (E): monoculture, E + S (CM): epithelial cells with 2D stromal cell conditioned medium, E + S (2L): 2-layer co-culture, N = 3, n = 3-5. One-way ANOVA, **P ≤ 0.01, *P ≤ 0.05.

### Multi-compartment assembloids mimic endometrial tissue-specific architecture and cellular composition

Using the methodology described in **Fig. 2a**, we seeded hEM3 (4 × 10^4^ cells/µL) in the assembloid core and 9700N (1 × 10^3^ cells/µL) in the corona to capture this ECM microenvironment *in vitro*. We used growth factor-reduced Matrigel (core), and 4 mg/mL collagen I (corona) in these assembloids. The conventional endometrial model (*41*) consists of embedding isolated epithelial cells in recombinant basement membrane (**Fig. 2b**), followed by a minimum of one week of culture in a specialized medium supplemented with growth factors and pathway inhibitors (*41*). This method supports the formation of hormone-responsive organoids and enables long-term expansion. However, the approach typically yields dozens to hundreds of organoids per well in bulk culture, resulting in significant heterogeneity in organoid size, morphology, and developmental state. These limitations, highlighted in subsequent studies and reviews (*34*, *44*), reduce experimental reproducibility and hinder physiological relevance due to lack of cell–cell and cell–matrix interactions and the absence of stromal–epithelial crosstalk. In contrast, our multi-compartment assembloids are uniform in size (*49*, *56*), owing to the spatial control enabled by compartmentalized cell and ECM seeding through oil-in-water droplet microtechnology. Each assembloid is individually isolated, eliminating organoid-organoid crosstalk and allowing consistent exposure to nutrients and signaling molecules. This system also enables medium-throughput production: we routinely generate multiple96-well plates, each containing a single assembloid per well, tailored to the specific requirements of the downstream experiments. Multiple plates can be fabricated in a single day, demonstrating the scalability and efficiency of this method for downstream applications.

**Figure 3.**
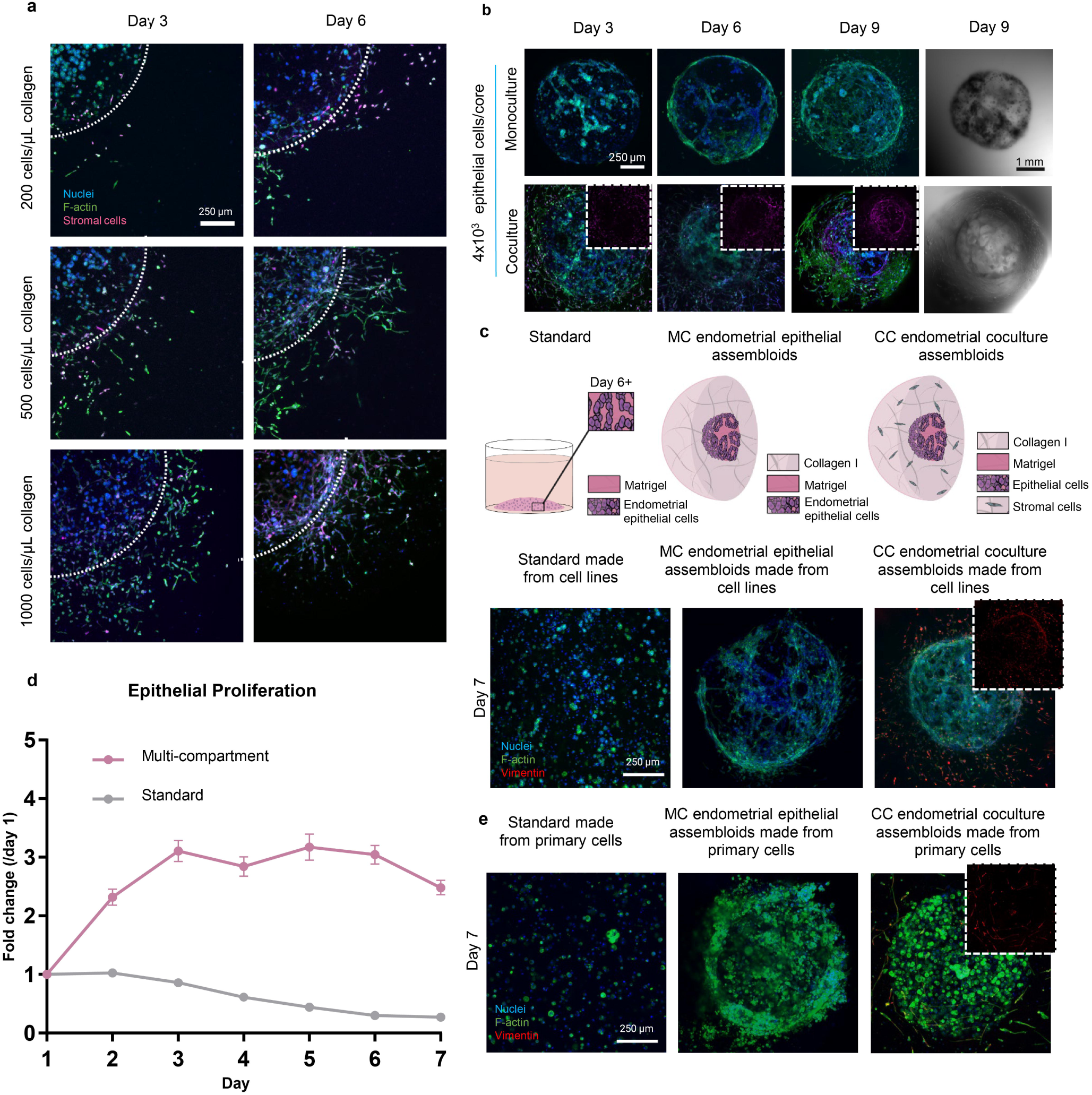
Multi-compartment endometrial assembloids mimic native tissue architecture and cell composition. (**A**) Development of the assembloid with low epithelial cell seeding density. Representative images of endometrial assembloid structure in the monoculture model seeded 1 × 10^4^ epithelial cells/core in coculture with 200, 500, and 1 × 10^3^ stromal cells/μL in the assembloid corona, from day 3 to day 9. Blue, nuclei. Green, F-actin. Magenta, stromal cells. Red, epithelial cells. For fluorescent images, scale bar, 250 µm (10X magnification). For phase contrast images, scale bar, 1 mm (4X magnification). (**B**) Development of the assembloid after raising epithelial cell seeding density. Representative images of endometrial assembloid structure in the monoculture model seeded at 4 × 10^4^ cells epithelial cells/core in coculture with gynecologic stromal cells in the assembloid corona, from day 3 to day 9. Blue, nuclei. Green, F-actin. Magenta, stromal cells. Red, epithelial cells. 10X magnification. For fluorescent images, scale bar, 250 µm; 10X magnification. For phase contrast images, scale bar, 300 µm; 10X magnification. (**C**) Comparison of the multi-compartment endometrial assembloid model with the standard organoid method, and representative fluorescent images of endometrial epithelial cells grown in the standard model (left), the monoculture (middle), and coculture (right) multi-compartment models at day 7. Blue, nuclei. Green, F-actin. Red, vimentin. Scale bar, 250 µm; 10X magnification. (**D**) PrestoBlue proliferation of total epithelial cells in both models. N=3, n=5+, data are mean ± SEM. Statistical test: t test, ns P > 0.05. Data are mean ± SEM. Significance is indicated as *p < 0.05, **p < 0.01, ***p < 0.001 ****p < 0.0001 (**E**) Immunofluorescence images of assembloid structures made from primary cells in the standard model, monoculture (MC) model seeded at 2 × 10^4^ epithelial cells/core, and in coculture (CC) with gynecologic stromal cells in the assembloid corona. Blue, nuclei. Green, F-actin. Red, stromal cells. 10X magnification. Scale bar, 250 µm.

For structural validation, the endometrial multi-compartment assembloids generated by our oil-in-water microtechnology were comparatively analyzed to conventional endometrial organoid models (**Fig 3c**). We tracked the epithelial proliferation for 7 consecutive days after generation using a Presto Blue proliferation assay (**Fig. 3d**). The cell proliferation pattern during epithelium formation in multi-compartment assembloids matched physiological expectations more closely than standard organoids, demonstrating cell proliferation at the first 3 days, followed by structural assembly during maturation (**Fig. 2e**). Endometrial epithelial cells have limited proliferation capacity, increasing in the first 3 days and staying consistent subsequently (**Fig. 3d**), but structural maturation occurs at or around Day 6 (**Fig. 2e**). Thus, we can conclude that the network structure is matured by cell reorganization and interaction with the ECMs during the later stages.

By comparing the epithelial structures of each model, we also identified an interlinked network in the multi-compartment model, which was not observed in the standard model (**Fig. 3c**). This organization more closely resembles the native architecture of the endometrium, as shown in the H&E-stained tissue section (**Fig. 1a**). This suggests that inclusion of the secondary stromal compartment not only provides appropriate cell–ECM type-specific interactions and paracrine signaling between epithelial and stromal cells, but also contributes to the establishment of a layered architecture that more closely resembles native endometrial tissue (Fig. 1). The multi-compartment endometrial assembloid architecture matures after culture for 3-6 days and can be maintained until at least Day 14 for extended experimental timelines. The structure of a mature assembloid with complex 3D interconnected epithelial networking in the Matrigel core is shown in **Fig. 3c**. This includes the scenario where the assembloid is cocultured with densely packed stromal cells in the collagen corona.

To more accurately replicate native endometrial physiology and establish a baseline for disease modeling, we generated assembloids using primary cells derived from endometrial tissue. Assembloids were made using the same parameters previously optimized for cell line–based assembloids: 4 × 10^4^ cells/µL epithelial cells in the Matrigel core, a stromal cell density of 1 × 10^3^ cells/μL, and a collagen I concentration of 4 mg/mL in the surrounding corona. The primary cells used were commercially sourced endometrial (uterine) epithelial cells (placed in the core) and normal human uterine fibroblasts (placed in the corona). Each cell type was cultured in the recommended medium provided by the supplier, as detailed in the Materials and Methods section. For coculture assembloids, primary reproductive epithelial cell medium was used to support optimal growth, given the sensitivity of epithelial cells under these conditions. Representative images of monoculture and coculture assembloids derived from primary cells are shown in **Fig. 3e**. Stromal cells in the coculture model were identified by vimentin immunostaining. Compared to established cell lines, primary endometrial cells are larger and require longer culture times, which extends the overall maturation process of the assembloid and results in a rounder epithelial cell morphology. However, they remain compatible with coculture conditions. Together, these findings confirm that our multi-compartment assembloid platform properly models the cellular architecture, growth dynamics, and tissue organization of the native endometrium.

### Multi-compartment assembloids reflect key molecular markers of native tissue

In native endometrial tissue, glandular epithelial cells express characteristic markers, including intermediate filament protein specific to epithelial cells cytokeratin 7 (CK7) and CK8, membrane-localized cell-cell adhesion protein E-cadherin which is essential for epithelial polarization and migration, estrogen receptor (ER), and epithelial cell adhesion molecule (EpCAM) (*41*, *56–59*). Accordingly, we performed immunofluorescence staining followed by confocal microscopy for CK8 and E-cadherin, which were both expressed in monoculture and coculture assembloids. The epithelial cells retained their standard organization and displayed continuous CK8 and E-cadherin staining (**Fig. 4a**), indicative of preserved epithelial identity and network formation. Western blot analysis showed that monoculture and coculture assembloids derived from cell lines (**Fig. 4b**) and primary cells (**Fig. 4c**) featured sustained expression of epithelial markers of CK7, CK8, EpCAM, and ER (*60*).

The stromal compartment expressed vimentin (**Fig. 3c,e**), confirming that each compartment maintained its lineage-specific marker profile and identity in both assembloids made either from cells lines or primary cells. To further assess epithelial organization, laminin distribution was analyzed as a marker of epithelial polarization. Laminin typically localizes to the stromal-facing basal surface of the epithelium (*43*). In monoculture assembloids, laminin staining revealed deposition in the basement membrane, confirming proper epithelial polarity within the assembloid architecture In monoculture assembloids, laminin staining revealed strong localization at the basolateral surface of epithelial cell clusters—specifically at the interface where the cells contact the Matrigel core—consistent with cell-deposited basement membrane and proper epithelial polarity. (**Fig. 4d**). These results demonstrate that our endometrial assembloids recapitulate key structural and molecular features of native epithelium, including lineage-specific marker expression, epithelial-stromal organization, and polarized architecture.

**Figure 4.**
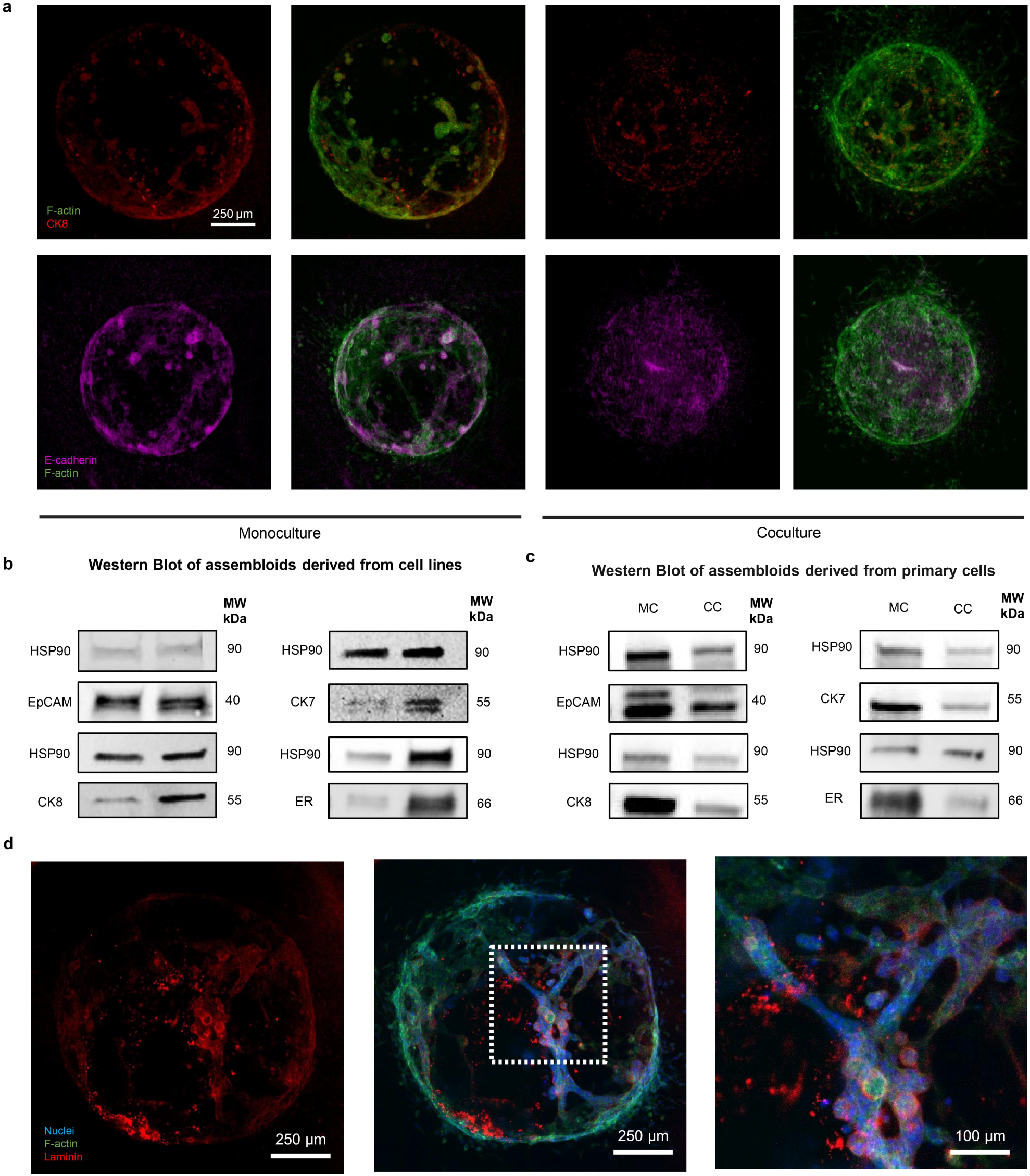
Multi-Compartment assembloids maintain key tissue-specific molecular markers. (**A**) Immunostaining of CK8 and E-cadherin in monoculture and coculture assembloids. CK8 (red), E-Cadherin (Magenta), and zoom-in overlay with F-actin. 10X magnification. (**B-C**) Western blot of assembloids derived from (**B**) cell lines and (**C**) primary cells to analyze the expression of epithelial markers EpCAM, cytokeratins 7 and 8, and Estrogen Receptor alpha; HSP90 expression was used as a loading control. (**D**) Immunostaining of laminin in the monoculture assembloids. Images from left to right are laminin (red), overlay with nuclei (blue) and F-actin (green), and zoom-in overlay with F-actin. 10X magnification.

### Multi-compartment assembloids recapitulate native tissue functionality and dynamic response

In the human endometrium, both epithelial and stromal cells respond to cyclic sex steroid hormones, with a series of changes regarding their morphological and molecular patterns (*61*). To observe whether our assembloids respond to the sex hormones, we designed a hormone treatment timeline (**Fig. 5a**). 17β-estradiol (E2) and progesterone (P4) were used to simulate the microenvironment of proliferative and secretory phases. RU-486 (also known as mifepristone) is a progesterone antagonist, which was used to simulate the menstrual phase. During the 2 weeks of treatment, phenol red-free culture medium was used, as phenol red is a weak estrogen (*62*).

Our results demonstrate a physiological response of our assembloid model to these hormone stimuli. Stromal cells experience molecular and morphological changes during the menstrual cycle. Decidualization is induced by P4 to prepare the endometrium for possible pregnancy (*63*), while the morphology of stromal cells changes from a spindle shape to a more rounded shape (*64*). We observed similar morphological changes in the stromal cells of the coculture assembloid models, from elongated to a circular shape after P4 stimulation (**Fig 5b**). This change in response to sex hormones indicated the possibility of decidualization in our endometrial assembloid, which was further confirmed via prolactin (PRL) relative expression (**Fig. 5h**) (*65*). Relative expression of progesterone receptor (PGR) in the coculture assembloid also changed in response to P4 presence. After P4 addition, the expression level of PGR was downregulated (**Fig. 5c**), as determined by the ratio of PGR expression to that of the control, which aligned with our expectations (*66*).

During the proliferative phase, rising levels of estrogen (E2) are known to induce PGR expression, particularly in the epithelial compartment (*67*, *68*). Once the cycle transitions into the secretory phase, progesterone (P4) levels increase, which induces a negative feedback loop that actively downregulates PR expression, especially in the glandular epithelium. This mechanism ensures that PR does not remain persistently active, preventing prolonged progesterone signaling which could disrupt endometrial function.

**Figure 5.**
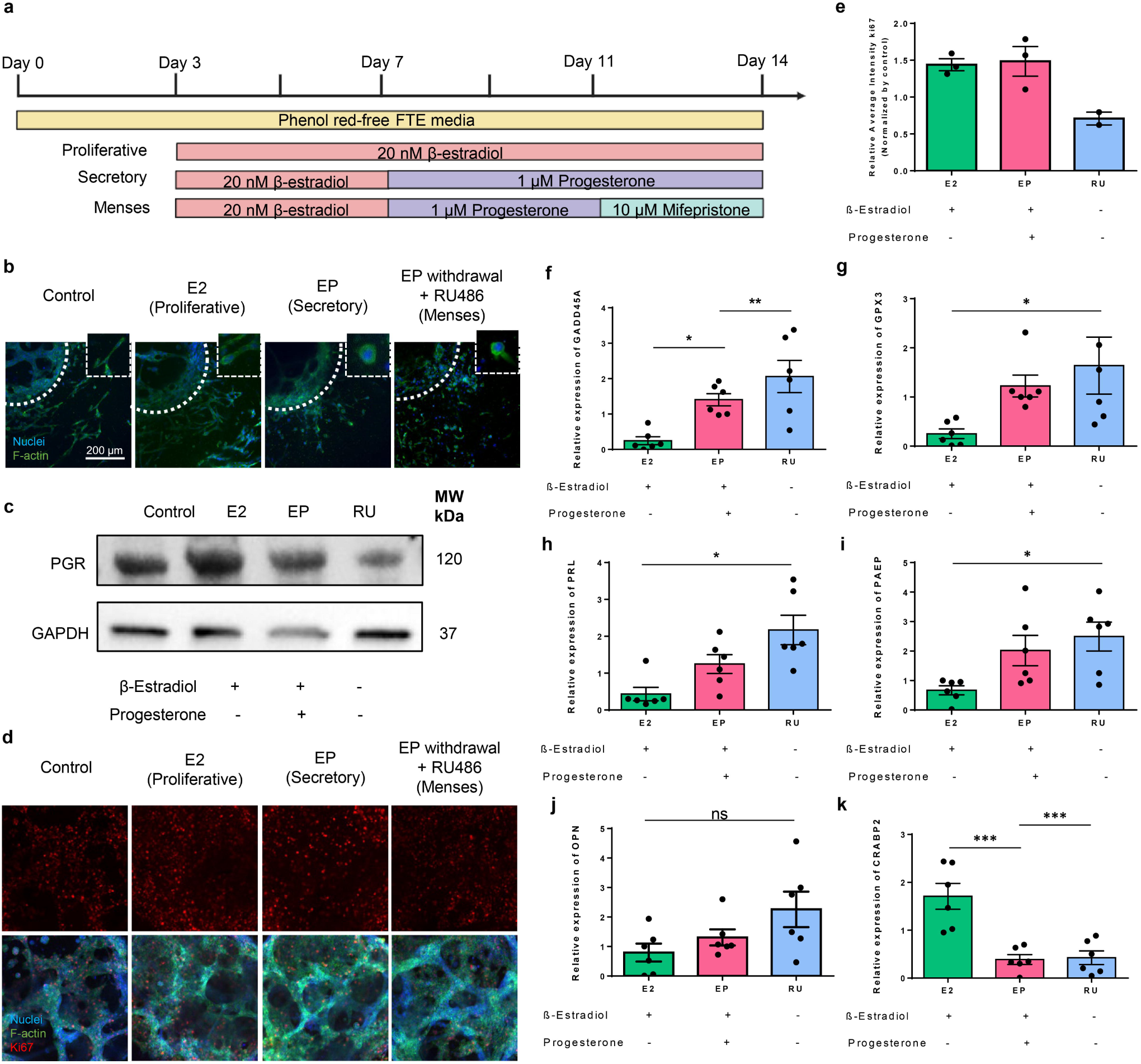
Hormonal regulation of morphology, proliferation, and gene expression in multi-compartment endometrial assembloids. (**A**) Hormone treatment timeline. Proliferation and apoptosis of endometrial assembloids in response to hormone stimulation. (**B**). Morphology of stromal cells in different treatment groups. Blue, nuclei. Green, F-actin. Scale bar = 200 µm; 20X magnification. (**C**). Normalized signal intensities of PGR to the loading control GAPDH was determined with image J analysis by normalizing the signal intensity of the target band divided the intensity of the loading control band. PGR/GAPDH: Control = 2.05, Proliferative phase = 3.47, Secretory phase = 2.72, Menses = 0.78. (**D**). Zoomed in Immunostaining of Ki67 (red) in coculture assembloid. Green, F-actin and Blue, Nuclei. Scale bar, 200 µm; 10X magnification. (**E**) Quantification of Ki67 expression as average fluorescence intensity per nuclei area. For each group, N = 3, n = 3 (**F-K)** RTqPCR Panel of gene response to hormonal treatment. (F) GADD45A, (G) GPX3, (H) PRL, (I) PAEP, (J) OPN, (K) CRABP2. For each group, N = 6, n = 3. **E** and **F**, Statistical test: one-way ANOVA, ns P > 0.05. Data are mean ± SEM. Significance is indicated as *p < 0.05, **p < 0.01, ***p < 0.001.

To capture cell proliferation, which is fundamental to the regenerative capacity of the endometrium, we used IF imaging to examine expression levels of the proliferation marker Ki67. After E2 was added, Ki67 was upregulated in the epithelium, which is characteristic of the proliferative phase (**Fig. 5d, e**). The withdrawal of E2 triggered a drop in the Ki67 expression in the IF, which also matched our expectations.

We measured the molecular response of the endometrial assembloids to the hormone treatment via RT-qPCR (**Fig. 5f-k**). Apoptosis marker Growth Arrest and DNA Damage Inducible alpha (GADD45A) helps regulate cell turnover in the endometrial epithelium and stroma, ensuring controlled proliferation (*69*, *70*). GADD45A significantly increased during the secretion phase as a result of P4 counter effect in proliferation, and there was once again another increase during the menses phase in response to RU-486, which induced apoptosis in the assembloid cells (**Fig. 5f**) Glutathione Peroxidase 3 (GPX3) acts as an antioxidant and plays a crucial role protecting the cells from damage, particularly during decidualization and implantation (*71*). PRL plays a role in preparing the endometrium for implantation and its expression coincides with decidualization, a process essential for implantation (*72*). Progesterone-associated endometrial protein (PAEP), also known as glycodelin, is involved in the establishment of endometrial receptivity for implantation (*73*, *74*). The expression of these three genes was upregulated with progesterone treatment, matching our expectations (**Fig. 5g-i**). Osteopontin (OPN) is a marker for stromal decidualization that mediates cell-cell and cell-matrix adhesion and is important for implantation and pregnancy success by facilitating blastocyst attachment to the endometrium (*75*, *76*). We observed a slight upregulation of this marker after the progesterone treatment (**Fig. 5j**), also, as expected. Cellular

Retinoic Acid Binding Protein 2 (CRABP2), a marker involved in tissue repair and regeneration expression after menstruation was reduced in our multi-compartment organoid upon progesterone treatment (**Fig. 5k**). This is consistent with in vivo biology, as P4 is known to suppress regenerative signaling during secretory phase (*77*).

Overall, these functional assessments demonstrate that our multi-compartment endometrial assembloids closely resemble the native endometrium in both the morphological and molecular responses to hormonal cues, confirming their functional similarity to the tissue and establishing them as a physiologically accurate platform for modeling endometrial dynamics across the menstrual cycle.

## Discussion

This study introduces a next-generation model of human physiology that unifies structural integrity, hormonal complexity, and fabrication scalability into a single, modular platform. We present an engineered endometrial assembloid model with compartmentalized cell and ECM regions resembling the tissue microanatomy revealed by a novel 3D reference map for human endometrial tissue. Endometrial epithelial cells are grown in a Matrigel core with stromal cells seeded in the collagen I corona, which precisely mimics the cell-cell and cell-ECM interactions of the tissue (*78*). The assembloid design parameters of each compartment were customized to structurally mimic *in situ* endometrial tissues. A core innovation of our platform is the oil column–based microfabrication method, which produces uniform, reproducible, and size-controlled assembloids using standard lab tools (*45*, *49*, *56*). These size-consistent assembloids allow spatial organization of ECM and cells to replicate realistic structural and biochemical cues (*79*). This approach enables long-term culture and avoids the need for synthetic hydrogel systems or bioprinting, making it accessible for a wide range of laboratories and ideal for mid-to-high-throughput experimental designs.

3D single-cell resolution CODA maps provide a framework for carefully selecting and optimizing the ECM compositions of each compartment based on histological data and carefully validating tissue architecture. The endometrial assembloids were validated to molecularly, functionally, and architecturally mimic human endometrium, and evidence was provided that these assembloids form interconnected epithelial networks and express lineage-specific markers. This was demonstrated in assembloids generated from primary cells and cell lines. Importantly, both the epithelial and stromal compartments respond appropriately to ovarian sex hormones, 17β-estradiol and progesterone, exhibiting expected morphological and molecular changes. Notably, stromal cells undergo decidualization upon hormone stimulation, a response that is uniquely captured in the multi-compartment configuration and not observed in standard organoid models. This highlights a key advantage of our model: the ability to dissect functional responses that are specific to one cell type yet depend on interactions with another cell type. Stromal cells and epithelial cells can be accurately cocultured in adjacent compartments where they maintain the cell-ECM and cell-cell signaling methods experienced in the *in situ* tissue and have independent functional responses to stimuli. This coculture also suggested that three is stromal regulation during epithelial development in addition to these independent functional responses. This provides a more accurate platform for studying epithelial-stromal interactions, which are essential for understanding normal tissue regeneration and disease progression.

Progesterone has been reported to antagonize Estrogen-induced epithelial proliferation (*80–82*), an effect that was not observed to the same extent in our assembloid model (**Fig. 5e**). One possible explanation is that the progesterone concentration used, based on established endometrial models (*83*), may not be optimal for the specific cell sources employed. Given that hormone responsiveness can vary significantly between primary and cultured cells or across donors, the lack of a proliferative shift could reflect cell-intrinsic differences in receptor expression or signaling thresholds. Adjusting the progesterone concentration or treatment duration may uncover additional regulatory effects. Notably, our model does exhibit robust hormone-induced stromal decidualization, highlighting a unique advantage of the multi-compartment design in capturing compartment-specific responses.

To our knowledge, this is the first study to evaluate the expression dynamics of OLFM1, EDN3, and MAOA across the full hormone cycle(*84*). OLFM1, a marker associated with extracellular matrix remodeling and implantation receptivity (*74*), EDN3, implicated in vascular signaling (*73*), and MAOA, associated with immune regulation and oxidative stress during the menstrual phase (*85*), remained largely unresponsive to hormonal cues (results not shown). These findings suggest that the regulation of these genes may depend on additional context-specific signals not present in our current model, such as interactions with trophoblasts, immune cells (e.g., NK cells or macrophages), or endothelial cells (*45*). Their lack of dynamic expression highlights both the specificity of our compartmentalized hormone responses and the importance of incorporating additional cellular populations in future iterations of the model to capture the full complexity of endometria.

Beyond physiological modeling, the system is designed with future extensibility in mind. It is readily adaptable for the integration of additional cell types, such as immune or endothelial cells or for coculture with trophoblast spheroids to model implantation (*45*). These features position the assembloid as a versatile tool for investigating not only normal endometrial biology, but also pathological conditions like endometriosis, recurrent implantation failure, and endometrial cancer. In sum, this multi-compartment assembloid platform offers a powerful and accessible system for investigating human endometrial structure and function across the menstrual cycle. Its architectural realism, dynamic hormone responsiveness, and translational potential provide a robust foundation for reproductive biology, disease modeling, and personalized medicine.

## Acknowledgments

The authors would like to thank all members of the Wirtz lab for their feedback. This work was partially supported through grants to D.W. from the National Cancer Institute (UG3CA275681, U54CA268083, and RO1CA300052). All cartoons were created with BioRender.com.

## Author contributions

K.R, A.C. and V.D developed the hypothesis and designed the experiments. V.D and K.R performed wet lab experiments and data analysis assisted by C.Z, B.H, G.M.A. The computational analysis was performed by A.F. Image analysis was performed by V.D and K.R. D.W. secured the funding for this work. The manuscript was written by V.D, K.R., and D.W. with input from C.Z, A.C, A.F, and B.H.

## Data and materials availability

All data are presented in the main or supplementary materials. All raw data and cell lines are available upon request.

## References

1. I.-S. Hong, Endometrial Stem Cells: Orchestrating Dynamic Regeneration of Endometrium and Their Implications in Diverse Endometrial Disorders. Int J Biol Sci 20, 864–879 (2024).

2. V. Makker, H. MacKay, I. Ray-Coquard, D. A. Levine, S. N. Westin, D. Aoki, A. Oaknin, Endometrial cancer. Nat Rev Dis Primers 7, 1–22 (2021).

3. K. T. Zondervan, C. M. Becker, K. Koga, S. A. Missmer, R. N. Taylor, P. Viganò, Endometriosis. Nat Rev Dis Primers 4, 1–25 (2018).

4. Endometriosis. https://www.who.int/news-room/fact-sheets/detail/endometriosis.

5. H. Mahdy, E. S. Vadakekut, D. Crotzer, “Endometrial Cancer” in StatPearls (StatPearls Publishing, Treasure Island (FL), 2025; http://www.ncbi.nlm.nih.gov/books/NBK525981/).

6. J. Evans, L. A. Salamonsen, A. Winship, E. Menkhorst, G. Nie, C. E. Gargett, E. Dimitriadis, Fertile ground: human endometrial programming and lessons in health and disease. Nat Rev Endocrinol 12, 654–667 (2016).

7. V. Makker, H. MacKay, I. Ray-Coquard, D. A. Levine, S. N. Westin, D. Aoki, A. Oaknin, Endometrial cancer. Nat Rev Dis Primers 7, 88 (2021).

8. W. B. Nothnick, C. Marsh, Z. Alali, Future Directions in Endometriosis Research and Therapeutics. Curr Womens Health Rev 14, 189–194 (2018).

9. L. Alzamil, K. Nikolakopoulou, M. Y. Turco, Organoid systems to study the human female reproductive tract and pregnancy. Cell Death Differ 28, 35–51 (2021).

10. A. R. Murphy, H. Campo, J. J. Kim, Strategies for modelling endometrial diseases. Nat Rev Endocrinol 18, 727–743 (2022).

11. N. Vogt, Assembloids. Nat Methods 18, 27–27 (2021).

12. J. Kim, B.-K. Koo, J. A. Knoblich, Human organoids: model systems for human biology and medicine. Nat Rev Mol Cell Biol 21, 571–584 (2020).

13. A. Forjaz, E. Vaz, V. M. Romero, S. Joshi, V. Queiroga, A. M. Braxton, A. C. Jiang, K. Fujikura, T. Cornish, S.-M. Hong, R. H. Hruban, P.-H. Wu, L. D. Wood, A. L. Kiemen, D. Wirtz, Three-dimensional assessments are necessary to determine the true, spatially resolved composition of tissues. Cell Reports Methods 5 (2025).

14. Long-term, hormone-responsive organoid cultures of human endometrium in a chemically defined medium | Nature Cell Biology. https://www.nature.com/articles/ncb3516.

15. J. S. Gnecco, A. Brown, K. Buttrey, C. Ives, B. A. Goods, L. Baugh, V. Hernandez-Gordillo, M. Loring, K. B. Isaacson, L. G. Griffith, Organoid co-culture model of the human endometrium in a fully synthetic extracellular matrix enables the study of epithelial-stromal crosstalk. Med 4, 554–579.e9 (2023).

16. T. M. Rawlings, K. Makwana, D. M. Taylor, M. A. Molè, K. J. Fishwick, M. Tryfonos, J. Odendaal, A. Hawkes, M. Zernicka-Goetz, G. M. Hartshorne, J. J. Brosens, E. S. Lucas, Modelling the impact of decidual senescence on embryo implantation in human endometrial assembloids. eLife 10, e69603 (2021).

17. A. Stejskalová, V. Fincke, M. Nowak, Y. Schmidt, K. Borrmann, M.-K. von Wahlde, S. D. Schäfer, L. Kiesel, B. Greve, M. Götte, Collagen I triggers directional migration, invasion and matrix remodeling of stroma cells in a 3D spheroid model of endometriosis. Sci Rep 11, 4115 (2021).

18. T. Wiwatpanit, A. R. Murphy, Z. Lu, M. Urbanek, J. E. Burdette, T. K. Woodruff, J. J. Kim, Scaffold-Free Endometrial Organoids Respond to Excess Androgens Associated With Polycystic Ovarian Syndrome. The Journal of Clinical Endocrinology & Metabolism 105, 769–780 (2020).

19. S. G. Zambuto, K. B. H. Clancy, B. A. C. Harley, A gelatin hydrogel to study endometrial angiogenesis and trophoblast invasion. Interface Focus 9, 20190016 (2019).

20. A. R. Murphy, T. Wiwatpanit, Z. Lu, B. Davaadelger, J. J. Kim, Generation of Multicellular Human Primary Endometrial Organoids. J Vis Exp, doi: 10.3791/60384 (2019).

21. E. Francés-Herrero, E. Juárez-Barber, H. Campo, S. López-Martínez, L. de Miguel-Gómez, A. Faus, A. Pellicer, H. Ferrero, I. Cervelló, Improved Models of Human Endometrial Organoids Based on Hydrogels from Decellularized Endometrium. Journal of Personalized Medicine 11, 504 (2021).

22. Y. Hibaoui, A. Feki, Organoid Models of Human Endometrial Development and Disease. Front Cell Dev Biol 8, 84 (2020).

23. K. Nikolakopoulou, M. Y. Turco, Investigation of infertility using endometrial organoids. doi: 10.1530/REP-20-0428 (2021).

24. S. De Vriendt, C. M. Casares, S. Rocha, H. Vankelecom, Matrix scaffolds for endometrium-derived organoid models. Front. Endocrinol. 14 (2023).

25. A. Luddi, V. Pavone, B. Semplici, L. Governini, M. Criscuoli, E. Paccagnini, M. Gentile, G. Morgante, V. De Leo, G. Belmonte, N. Zarovni, P. Piomboni, Organoids of Human Endometrium: A Powerful In Vitro Model for the Endometrium-Embryo Cross-Talk at the Implantation Site. Cells 9, 1121 (2020).

26. X. Jiang, X. Li, X. Fei, J. Shen, J. Chen, M. Guo, Y. Li, Endometrial membrane organoids from human embryonic stem cell combined with the 3D Matrigel for endometrium regeneration in asherman syndrome. Bioactive Materials 6, 3935–3946 (2021).

27. T. M. Rawlings, K. Makwana, M. Tryfonos, E. S. Lucas, Organoids to model the endometrium: implantation and beyond. doi: 10.1530/RAF-21-0023 (2021).

28. S.-R. Park, S.-R. Kim, J. B. Im, C. H. Park, H.-Y. Lee, I.-S. Hong, 3D stem cell-laden artificial endometrium: successful endometrial regeneration and pregnancy. Biofabrication 13, 045012 (2021).

29. J. Ahn, M.-J. Yoon, S.-H. Hong, H. Cha, D. Lee, H. S. Koo, J.-E. Ko, J. Lee, S. Oh, N. L. Jeon, Y.-J. Kang, Three-dimensional microengineered vascularised endometrium-on-a-chip. Human Reproduction 36, 2720– 2731 (2021).

30. H. Campo, A. Murphy, S. Yildiz, T. Woodruff, I. Cervelló, J. J. Kim, Microphysiological Modeling of the Human Endometrium. Tissue Eng Part A 26, 759–768 (2020).

31. S. Muruganandan, X. Fan, S. Dhal, N. R. Nayak, Development of A 3D Tissue Slice Culture Model for the Study of Human Endometrial Repair and Regeneration. Biomolecules 10, 136 (2020).

32. N. Nie, L. Gong, D. Jiang, Y. Liu, J. Zhang, J. Xu, X. Yao, B. Wu, Y. Li, X. Zou, 3D bio-printed endometrial construct restores the full-thickness morphology and fertility of injured uterine endometrium. Acta Biomaterialia 157, 187–199 (2023).

33. M. Yamaguchi, K. Yoshihara, N. Yachida, K. Suda, R. Tamura, T. Ishiguro, T. Enomoto, The New Era of Three-Dimensional Histoarchitecture of the Human Endometrium. Journal of Personalized Medicine 11, 713 (2021).

34. H. C. Fitzgerald, D. J. Schust, T. E. Spencer, In vitro models of the human endometrium: evolution and application for women’s health. Biol Reprod 104, 282–293 (2021).

35. X. Li, S. P. Kodithuwakku, R. W. S. Chan, W. S. B. Yeung, Y. Yao, E. H. Y. Ng, P. C. N. Chiu, C.-L. Lee, Three-dimensional culture models of human endometrium for studying trophoblast-endometrium interaction during implantation. Reproductive Biology and Endocrinology 20, 120 (2022).

36. C. M. Klapperich, G. Vunjak-Novakovic, J. Robinson, R. Horton, P. K. Kreeger, M. Mahmoudi, S. Shah, A. Frolova, Bioengineering solutions to improve women’s health. Med 6 (2025).

37. Y. Abbas, L. G. Brunel, M. S. Hollinshead, R. C. Fernando, L. Gardner, I. Duncan, A. Moffett, S. Best, M. Y. Turco, G. J. Burton, R. E. Cameron, Generation of a three-dimensional collagen scaffold-based model of the human endometrium. Interface Focus 10, 20190079 (2020).

38. A. D. van den Brand, E. Rubinstein, P. C. de Jong, M. van den Berg, M. B. M. van Duursen, Primary endometrial 3D co-cultures: A comparison between human and rat endometrium. The Journal of Steroid Biochemistry and Molecular Biology 194, 105458 (2019).

39. J. R. H. Wendel, X. Wang, L. J. Smith, S. M. Hawkins, Three-Dimensional Biofabrication Models of Endometriosis and the Endometriotic Microenvironment. Biomedicines 8, 525 (2020).

40. H. Wang, F. Pilla, S. Anderson, S. Martínez-Escribano, I. Herrer, J. M. Moreno-Moya, S. Musti, S. Bocca, S. Oehninger, J. A. Horcajadas, A novel model of human implantation: 3D endometrium-like culture system to study attachment of human trophoblast (Jar) cell spheroids. Mol Hum Reprod 18, 33–43 (2012).

41. M. Y. Turco, L. Gardner, J. Hughes, T. Cindrova-Davies, M. J. Gomez, L. Farrell, M. Hollinshead, S. G. E. Marsh, J. J. Brosens, H. O. Critchley, B. D. Simons, M. Hemberger, B.-K. Koo, A. Moffett, G. J. Burton, Long-term, hormone-responsive organoid cultures of human endometrium in a chemically defined medium. Nat Cell Biol 19, 568–577 (2017).

42. M. Boretto, B. Cox, M. Noben, N. Hendriks, A. Fassbender, H. Roose, F. Amant, D. Timmerman, C. Tomassetti, A. Vanhie, C. Meuleman, M. Ferrante, H. Vankelecom, Development of organoids from mouse and human endometrium showing endometrial epithelium physiology and long-term expandability. Development 144, 1775–1786 (2017).

43. J. S. Gnecco, A. Brown, K. Buttrey, C. Ives, B. A. Goods, L. Baugh, V. Hernandez-Gordillo, M. Loring, K. B. Isaacson, L. G. Griffith, Organoid co-culture model of the human endometrium in a fully synthetic extracellular matrix enables the study of epithelial-stromal crosstalk. Med 4, 554–579.e9 (2023).

44. T. M. Rawlings, K. Makwana, M. Tryfonos, E. S. Lucas, Organoids to model the endometrium: implantation and beyond. Reprod Fertil 2, R85–R101 (2021).

45. M.-H. Lee, G. C. Russo, Y. S. Rahmanto, W. Du, A. J. Crawford, P.-H. Wu, D. Gilkes, A. Kiemen, T. Miyamoto, Y. Yu, M. Habibi, I.-M. Shih, T.-L. Wang, D. Wirtz, Multi-compartment tumor organoids. Materials Today 61, 104–116 (2022).

46. S. Joshi, A. Forjaz, K. S. Han, Y. Shen, V. Queiroga, F. A. Selaru, M. Gérard, D. Xenes, J. Matelsky, B. Wester, A. M. Barrutia, A. L. Kiemen, P.-H. Wu, D. Wirtz, InterpolAI: deep learning-based optical flow interpolation and restoration of biomedical images for improved 3D tissue mapping. Nat Methods, 1–12 (2025).

47. A. M. Braxton, A. L. Kiemen, M. P. Grahn, A. Forjaz, J. Parksong, J. Mahesh Babu, J. Lai, L. Zheng, N. Niknafs, L. Jiang, H. Cheng, Q. Song, R. Reichel, S. Graham, A. I. Damanakis, C. G. Fischer, S. Mou, C. Metz, J. Granger, X.-D. Liu, N. Bachmann, Y. Zhu, Y. Liu, C. Almagro-Pérez, A. C. Jiang, J. Yoo, B. Kim, S. Du, E. Foster, J. Y. Hsu, P. A. Rivera, L. C. Chu, F. Liu, E. K. Fishman, A. Yuille, N. J. Roberts, E. D. Thompson, R. B. Scharpf, T. C. Cornish, Y. Jiao, R. Karchin, R. H. Hruban, P.-H. Wu, D. Wirtz, L. D. Wood, 3D genomic mapping reveals multifocality of human pancreatic precancers. Nature 629, 679–687 (2024).

48. A. L. Kiemen, A. M. Braxton, M. P. Grahn, K. S. Han, J. M. Babu, R. Reichel, A. C. Jiang, B. Kim, J. Hsu, F. Amoa, S. Reddy, S.-M. Hong, T. C. Cornish, E. D. Thompson, P. Huang, L. D. Wood, R. H. Hruban, D. Wirtz, P.-H. Wu, CODA: quantitative 3D reconstruction of large tissues at cellular resolution. Nat Methods 19, 1490– 1499 (2022).

49. A. J. Crawford, A. Forjaz, J. Bons, I. Bhorkar, T. Roy, D. Schell, V. Queiroga, K. Ren, D. Kramer, W. Huang, G. C. Russo, M.-H. Lee, P.-H. Wu, I.-M. Shih, T.-L. Wang, M. A. Atkinson, B. Schilling, A. L. Kiemen, D. Wirtz, Combined assembloid modeling and 3D whole-organ mapping captures the microanatomy and function of the human fallopian tube. Science Advances 10, eadp6285 (2024).

50. A. Forjaz, V. M. Romero, I. Reucroft, M. Eminizer, D. Kramer, D. Higuera, H. Mojdeganlou, P. A. Guerrero, J. Min, M. Wetzel, D. Lvovs, A. Valentin, S. M. Shin, X. Yuan, R. C. Sears, K. Chin, A. Maitra, E. J. Fertig, W. J. Ho, L. T. Kagohara, L. D. Wood, D. Wirtz, D. N. Sidiropoulos, A. L. Kiemen, PIVOT: an open-source tool for multi-omic spatial data registration. bioRxiv [Preprint] (2025). 10.1101/2025.06.08.658506.

51. J. F. Dekkers, M. Alieva, L. M. Wellens, H. C. R. Ariese, P. R. Jamieson, A. M. Vonk, G. D. Amatngalim, H. Hu, K. C. Oost, H. J. G. Snippert, J. M. Beekman, E. J. Wehrens, J. E. Visvader, H. Clevers, A. C. Rios, High-resolution 3D imaging of fixed and cleared organoids. Nat Protoc 14, 1756–1771 (2019).

52. L. A. Salamonsen, J. C. Hutchison, C. E. Gargett, Cyclical endometrial repair and regeneration. Development 148, dev199577 (2021).

53. C. D. Cook, A. S. Hill, M. Guo, L. Stockdale, J. P. Papps, K. B. Isaacson, D. A. Lauffenburger, L. G. Griffith, Local remodeling of synthetic extracellular matrix microenvironments by co-cultured endometrial epithelial and stromal cells enables long-term dynamic physiological function. Integrative Biology 9, 271–289 (2017).

54. A. Stejskalová, V. Fincke, M. Nowak, Y. Schmidt, K. Borrmann, M.-K. von Wahlde, S. D. Schäfer, L. Kiesel, B. Greve, M. Götte, Collagen I triggers directional migration, invasion and matrix remodeling of stroma cells in a 3D spheroid model of endometriosis. Sci Rep 11, 4115 (2021).

55. J. T. Arnold, D. G. Kaufman, M. Seppälä, B. A. Lessey, Endometrial stromal cells regulate epithelial cell growth in vitro: a new co-culture model. Human Reproduction 16, 836–845 (2001).

56. G. C. Russo, A. J. Crawford, D. Clark, J. Cui, R. Carney, M. N. Karl, B. Su, B. Starich, T.-S. Lih, P. Kamat, Q. Zhang, P. R. Nair, P.-H. Wu, M.-H. Lee, H. S. Leong, H. Zhang, V. W. Rebecca, D. Wirtz, E-cadherin interacts with EGFR resulting in hyper-activation of ERK in multiple models of breast cancer. Oncogene 43, 1445–1462 (2024).

57. Y. Suryo Rahmanto, W. Shen, X. Shi, X. Chen, Y. Yu, Z.-C. Yu, T. Miyamoto, M.-H. Lee, V. Singh, R. Asaka, G. Shimberg, M. I. Vitolo, S. S. Martin, D. Wirtz, R. Drapkin, J. Xuan, T.-L. Wang, I.-M. Shih, Inactivation of Arid1a in the endometrium is associated with endometrioid tumorigenesis through transcriptional reprogramming. Nat Commun 11, 2717 (2020).

58. C. Sancakli Usta, G. Turan, C. B. Bulbul, A. Usta, E. Adali, Differential expression of Oct-4, CD44, and E-cadherin in eutopic and ectopic endometrium in ovarian endometriomas and their correlations with clinicopathological variables. Reprod Biol Endocrinol 18, 116 (2020).

59. S. Bajpai, J. Correia, Y. Feng, J. Figueiredo, S. X. Sun, G. D. Longmore, G. Suriano, D. Wirtz, α-Catenin mediates initial E-cadherin-dependent cell–cell recognition and subsequent bond strengthening. Proceedings of the National Academy of Sciences 105, 18331–18336 (2008).

60. M. Trzpis, P. M. J. McLaughlin, L. M. F. H. de Leij, M. C. Harmsen, Epithelial Cell Adhesion Molecule: More than a Carcinoma Marker and Adhesion Molecule. The American Journal of Pathology 171, 386–395 (2007).

61. N. Maenhoudt, A. De Moor, H. Vankelecom, Modeling Endometrium Biology and Disease. J Pers Med 12, 1048 (2022).

62. Y. Berthois, J. A. Katzenellenbogen, B. S. Katzenellenbogen, Phenol red in tissue culture media is a weak estrogen: implications concerning the study of estrogen-responsive cells in culture. Proceedings of the National Academy of Sciences 83, 2496–2500 (1986).

63. H. Okada, T. Tsuzuki, H. Murata, Decidualization of the human endometrium. Reproductive Medicine and Biology 17, 220–227 (2018).

64. B. Pan-Castillo, S. A. Gazze, S. Thomas, C. Lucas, L. Margarit, D. Gonzalez, L. W. Francis, R. S. Conlan, Morphophysical dynamics of human endometrial cells during decidualization. *Nanomedicine: Nanotechnology*, Biology and Medicine 14, 2235–2245 (2018).

65. H. Okada, T. Tsuzuki, H. Murata, Decidualization of the human endometrium. Reprod Med Biol 17, 220–227 (2018).

66. M. J. Large, F. J. DeMayo, The regulation of embryo implantation and endometrial decidualization by progesterone receptor signaling. Molecular and Cellular Endocrinology 358, 155–165 (2012).

67. J. D. Graham, C. L. Clarke, Physiological Action of Progesterone in Target Tissues*. Endocrine Reviews 18, 502–519 (1997).

68. M. Van Wynendaele, C. Thieffry, L. Samain, C. E. Pierreux, D. Tyteca, E. Marbaix, P. Henriet, Effects of estradiol, progesterone or cAMP on expression of PGRMC1 and progesterone receptor in a xenograft model of human endometrium and in endometrial cell culture. Steroids 198, 109284 (2023).

69. Gadd45a growth arrest and DNA-damage-inducible 45 alpha [Mus musculus (house mouse)] - Gene - NCBI. https://www.ncbi.nlm.nih.gov/gene/13197.

70. L. T. Kaufmann, C. Niehrs, *Gadd45a* and *Gadd45g* regulate neural development and exit from pluripotency in *Xenopus*. Mechanisms of Development 128, 401–411 (2011).

71. X. Xu, J.-Y. Leng, F. Gao, Z.-A. Zhao, W.-B. Deng, X.-H. Liang, Y.-J. Zhang, Z.-R. Zhang, M. Li, A.-G. Sha, Z.-M. Yang, Differential expression and anti-oxidant function of glutathione peroxidase 3 in mouse uterus during decidualization. FEBS Lett 588, 1580–1589 (2014).

72. R. L. Jones, H. O. Critchley, J. Brooks, H. N. Jabbour, A. S. McNeilly, Localization and temporal expression of prolactin receptor in human endometrium. J Clin Endocrinol Metab 83, 258–262 (1998).

73. L. C. Hautala, P.-C. Pang, A. Antonopoulos, A. Pasanen, C.-L. Lee, P. C. N. Chiu, W. S. B. Yeung, M. Loukovaara, R. Bützow, S. M. Haslam, A. Dell, H. Koistinen, Altered glycosylation of glycodelin in endometrial carcinoma. Lab Invest 100, 1014–1025 (2020).

74. J. E. Pearson-Farr, G. Wheway, M. S. A. Jongen, P. Goggin, R. M. Lewis, Y. Cheong, J. K. Cleal, Endometrial gland-specific progestagen-associated endometrial protein and cilia gene splicing changes in recurrent pregnancy loss. Reprod Fertil 3, 162–172 (2022).

75. G. A. Johnson, R. C. Burghardt, F. W. Bazer, T. E. Spencer, Osteopontin: Roles in Implantation and Placentation1. Biology of Reproduction 69, 1458–1471 (2003).

76. G. A. Johnson, R. C. Burghardt, F. W. Bazer, Osteopontin: a leading candidate adhesion molecule for implantation in pigs and sheep. Journal of Animal Science and Biotechnology 5, 56 (2014).

77. Q. Yang, R. Wang, W. Xiao, F. Sun, H. Yuan, Q. Pan, Cellular Retinoic Acid Binding Protein 2 Is Strikingly Downregulated in Human Esophageal Squamous Cell Carcinoma and Functions as a Tumor Suppressor. PLOS ONE 11, e0148381 (2016).

78. Physiology of the Endometrium and Regulation of Menstruation. 10.1152/physrev.00031.2019.

79. 79. A. J. Crawford, A. Johnston, W. Du, E. A. Hanna, D. Schell, Z. Wan, T.-H. Chen, F. Wu, K. Ren, Y. Lim, P. Nair, D. Wirtz, A 3D in vitro assay to study combined immune cell infiltration and cytotoxicity. bioRxiv [Preprint] (2024). 10.1101/2024.03.27.586980.

80. M. Y. Turco, L. Gardner, J. Hughes, T. Cindrova-Davies, M. J. Gomez, L. Farrell, M. Hollinshead, S. G. E. Marsh, J. J. Brosens, H. O. Critchley, B. D. Simons, M. Hemberger, B.-K. Koo, A. Moffett, G. J. Burton, Long-term, hormone-responsive organoid cultures of human endometrium in a chemically defined medium. Nat Cell Biol 19, 568–577 (2017).

81. L. Deligdisch, Hormonal Pathology of the Endometrium. Modern Pathology 13, 285–294 (2000).

82. H. L. Franco, C. A. Rubel, M. J. Large, M. Wetendorf, R. Fernandez-Valdivia, J.-W. Jeong, T. E. Spencer, R. R. Behringer, J. P. Lydon, F. J. DeMayo, Epithelial progesterone receptor exhibits pleiotropic roles in uterine development and function. FASEB J 26, 1218–1227 (2012).

83. H. C. Fitzgerald, P. Dhakal, S. K. Behura, D. J. Schust, T. E. Spencer, Self-renewing endometrial epithelial organoids of the human uterus. Proceedings of the National Academy of Sciences 116, 23132–23142 (2019).

84. S. Altmäe, M. Koel, U. Võsa, P. Adler, M. Suhorutšenko, T. Laisk-Podar, V. Kukushkina, M. Saare, A. Velthut-Meikas, K. Krjutškov, L. Aghajanova, P. G. Lalitkumar, K. Gemzell-Danielsson, L. Giudice, C. Simón, A. Salumets, Meta-signature of human endometrial receptivity: a meta-analysis and validation study of transcriptomic biomarkers. Sci Rep 7, 10077 (2017).

85. Z. Yu, P. Huang, L. Wang, F. Meng, Q. Shi, X. Huang, L. Qiu, H. Wang, S. Kong, J. Wu, Monoamine oxidases activity maintains endometrial monoamine homeostasis and participates in embryo implantation and development. BMC Biology 22, 166 (2024).

